# Comprehensive analysis of lung cancer pathology images to discover tumor shape features that predict survival outcome

**DOI:** 10.1101/274332

**Authors:** Shidan Wang, Alyssa Chen, Lin Yang, Ling Cai, Yang Xie, Junya Fujimoto, Adi Gazdar, Guanghua Xiao

**Affiliations:** Quantitative Biomedical Research Center, Department of Clinical Sciences, University of Texas Southwestern Medical Center, Dallas, Texas, 75390, USA; Department of Computer Sciences, Massachusetts Institute of Technology; Department of Pathology, National Cancer Center/Cancer Hospital, Chinese Academy of Medical Sciences and Peking Union Medical College, Beijing, 100021, China; Children’s Medical Center Research Institute at UT Southwestern Medical Center, 5323 Harry Hines Blvd, Dallas, TX, 75390 USA; Department of Bioinformatics, UT Southwestern Medical Center, Dallas, Texas, USA; Simmons Comprehensive Cancer Center, UT Southwestern Medical Center, Dallas, Texas, USA; Department of Translational Molecular Pathology, Division of Pathology/Lab Medicine, University of Texas MD Anderson Cancer Center, Houston, TX; Department of Pathology, University of Texas Southwestern Medical Center, Dallas, Texas, 75390, USA; Hamon Center for Therapeutic Oncology Research, UT Southwestern Medical Center, Texas, 75390, USA

## Abstract

Pathology slide images capture tumor histomorphological details in high resolution. However, manual detection and characterization of tumor regions in pathology slides is labor intensive and subjective. Using a deep convolutional neural network (CNN), we developed an automated tumor region recognition system for lung cancer pathology slides. From the identified regions, we extracted 22 well-defined tumor shape features and found that 15 of them were significantly associated with patient survival outcome in lung adenocarcinoma patients from the National Lung Screening Trial. A tumor shape-based prognostic model was developed and validated in an independent patient cohort (n=389). The predicted high-risk group had significantly worse survival than the low-risk group (p value = 0.0029). Predicted risk group serves as an independent prognostic factor (high-risk vs. low-risk, hazard ratio = 2.25, 95% CI 1.34-3.77, p value = 0.0022) after adjusting for age, gender, smoking status, and stage. This study provides new insights into the relationship between tumor shape and patient prognosis.

---

Lung cancer is the leading cause of death from cancer, with about half of all cases comprised of lung adenocarcinoma (ADC), which is remarkably heterogeneous in morphological features^1,2^ and highly variable in prognosis. Through sophisticated visual inspection of tumor pathology slides, ADC can be further classified into different subtypes with drastically different prognoses. Some contributing morphological features have been recognized, such as tumor size or vascular invasion in lung ADC. However, there is a lack of systematic studies on the relationship between tumor shape in pathology slides and patient prognosis.

Tumor tissue slide scanning is becoming part of routine clinical practice for the acquisition of high resolution tumor histological details. In recent years, several computer algorithms for hematoxylin and eosin (H&E) stained pathology image analysis have been developed to aid pathologists in objective clinical diagnosis and prognosis^3–7^. Examples include an algorithm to extract stromal features^8^ and an algorithm to assess cellular heterogeneity^6^ as a prognostic factor in breast cancer. More recently, studies have shown that morphological features are associated with patient prognosis in lung cancer as well^4,5,7^. Deep learning methods, such as convolution neural networks (CNNs), have been widely used in image segmentation, object classification and recognition^9–11^ and are now being adapted in biomedical image analysis to facilitate cancer diagnosis. To some extent, the performances of deep learning algorithms are similar to, or sometimes even better than, those of humans^12,13^. For analysis of H&E-stained pathology images, deep learning methods have been developed to distinguish tumor regions^14^, detect metastasis^15^, predict mutation status^16^, and classify tumors^17^ in breast cancer as well as in other cancers. However, due to the complexity of lung cancer tissue structures (such as microscopic alveoli and micro-vessel), deep learning methods for automatic lung cancer region detection from H&E-stained pathology images are not currently available.

Automatic tumor region detection allows for tumor size calculation and tumor shape estimation. Tumor size is a well-established lung cancer prognostic factor^18^; the effect of tumor shape has also been investigated in regard to its relationship with drug delivery^19,20^ and prognosis prediction^21–26^. In X-Ray and computer tomography (CT) image studies, the rough tumor boundary has been reported as a marker for malignant tumor in breast cancer^27^, and found to be associated with local tumor progression and worse prognosis in lung cancer patients^22,26^. Compared with CT images, which are most commonly used to evaluate tumor shape, pathology images have much higher spatial resolution^28^. Thus, automatic tumor region detection in pathology images allows us to characterize tumor shape accurately and extract tumor shape-based features.

In this study, we developed a deep CNN model to automatically recognize tumor regions for lung ADC from H&E pathology images. More importantly, based on a systematic study of the detected tumor regions of lung ADC patients from the National Lung Screening Trial (NLST) cohort (n=150), we found that many features that characterize the shape of the tumors are significantly associated with tumor prognosis. Finally, we developed a risk-prediction model for lung cancer prognosis using the tumor shape-related features from the NLST lung ADC patient cohort. The prognostic model was then validated in lung ADC patients from The Cancer Genome Atlas (TCGA) dataset. The design of this study is summarized in a flow chart (Figure 1).

**Figure 1.**
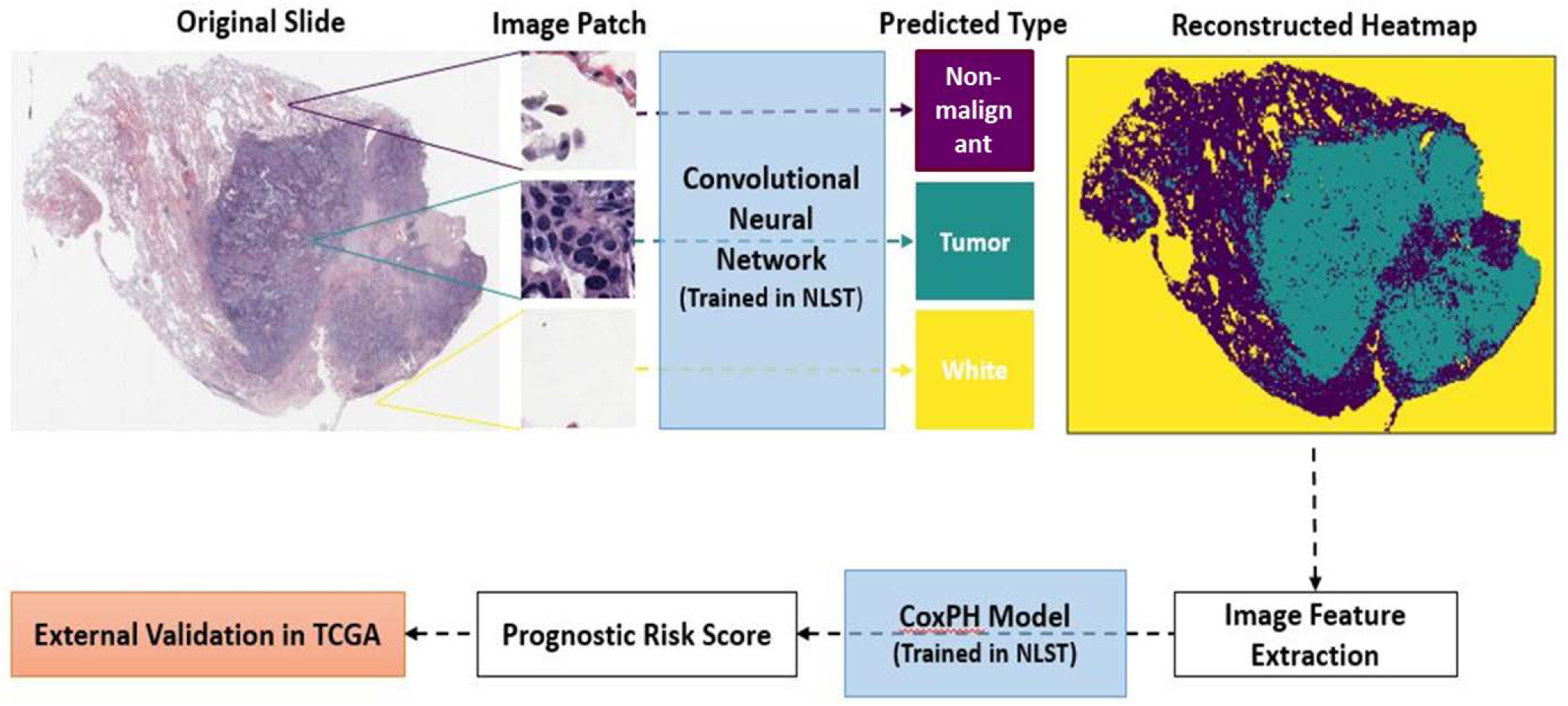
Flow chart of analysis process. CNN, convolutional neural network; NLST, the National Lung Screening Trial; TCGA, The Cancer Genome Atlas.

## RESULTS

### CNN model distinguishes tumor patches from non-malignant and white (empty region) patches

5344 tumor, non-malignant, and white image patches were extracted from 27 lung ADC H&E pathology slide images (Supplemental Figure 1). The imaging patches were split into training, validation and testing datasets (see Methods Section). The CNN model was trained on the training set. The training process stopped at the 28^th^ epoch after validation accuracy failed to improve after 10 epochs. The learning curves for the CNN model in the training and validation sets are shown in Supplemental Figure 2. The overall prediction accuracy of the CNN model in the testing set was 89.8%; the accuracy was 88.1% for tumor patches and 93.5% for non-malignant patches (Supplemental Table 1).

### Tumor region recognition for pathology images

In the NLST dataset, the pathology images have sizes ranging from 5280 × 4459 pixels to 36966 × 22344 pixels (median 24244 × 19261 pixels). To identify tumor regions at slide level, each whole slide image was partitioned into 300×300 image patches. To speed up slide-level prediction, tissue regions were first identified (see Methods Section) and only the image patches within the tissue regions were predicted by the CNN model (Supplemental Figure 3). The predicted probabilities of the image patches were summarized into heatmaps of tumor probability (Figure 2). An example of a tumor probability heatmap is shown in Figure 2B. The tumor region heatmap, predicted as the category with highest probability, is shown in Figure 2C. Each pixel in the heatmaps corresponds to a 300 × 300 pixel image patch in the original 40X pathology image.

**Figure 2.**
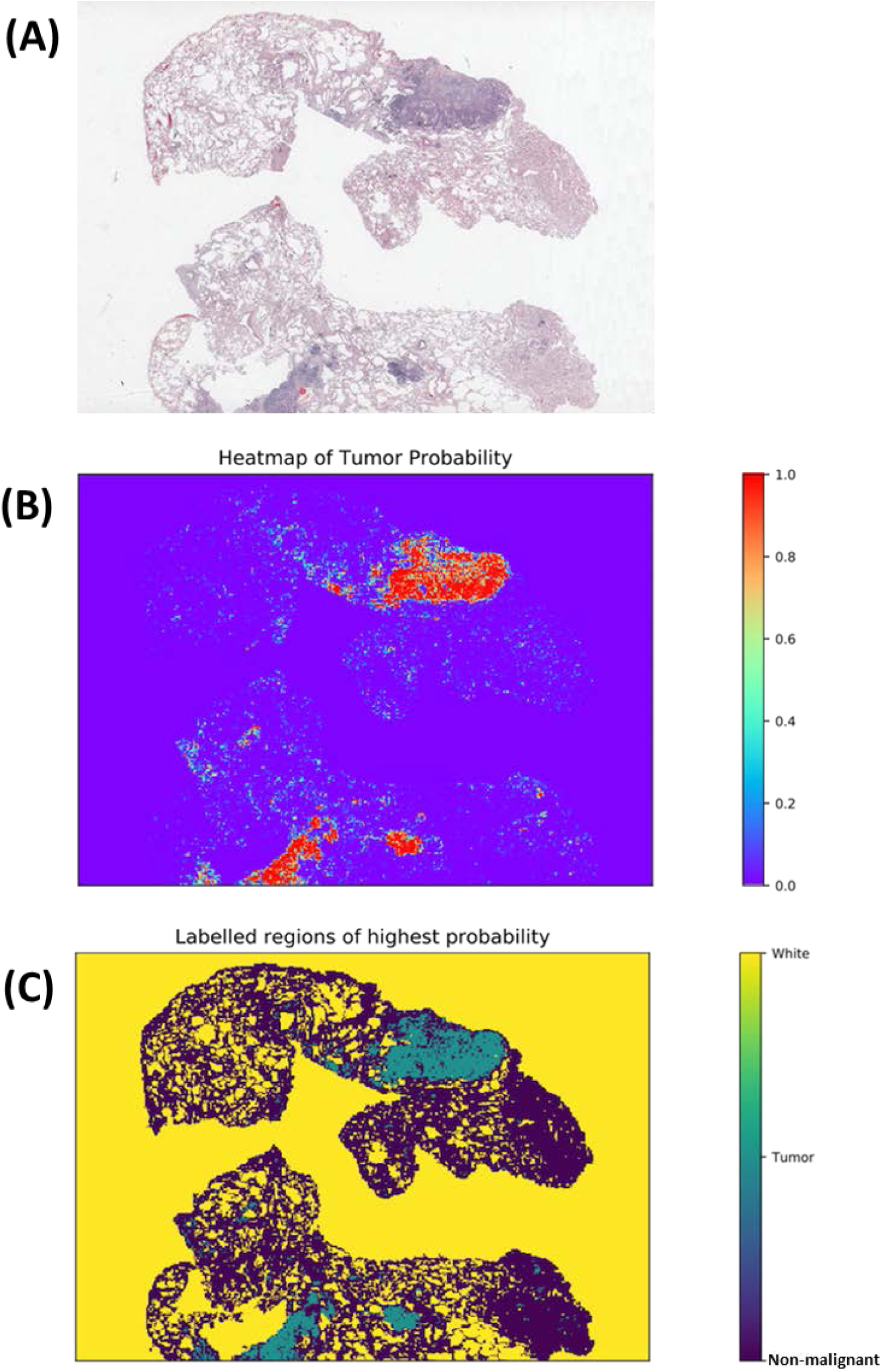
Example results of slide-level tumor region detection. (A) Original slide. (B) Predicted tumor probability. Each point in the heatmap corresponds to a 300 × 300 pixel image patch in original 40x slide. (C) Predicted region labels. Yellow: white (empty) region; green: tumor region; blue: non-malignant region.

### Image features from predicted tumor regions correlate with survival outcome

Based on the predicted tumor region heatmap, tissue samples were identified (Supplemental Figure 4) and 22 image features were extracted for each tissue sample (see Methods Section). For each patient the image features from multiple tissue samples of the same patient were averaged. The associations between tumor region features and prognostic outcome are summarized in Table 2 in the NLST dataset. It shows that many features were associated with survival outcome. Most tumor area/size-related features, including area, perimeter, convex area, filled area, major axis length, and minor axis length, both for all tumor regions and for the main tumor region, were associated with poor survival outcome. Interestingly, the number of holes and the perimeter^2 to area ratio, an estimation of circularity, were also associated with poor survival outcome (for all tumor regions: per 100 number of holes, hazard ratio [HR] = 1.087, p value = 0.033; per 1000 perimeter^2 to area ratio, HR = 1.15, p value = 0.016; similar results for main tumor region; Table 2). Examples comparing tumor shapes with high and low values of eccentricity and perimeter^2 to area ratio of main tumor region are illustrated in Figure 3. As expected, the angle between the X-axis and the major axis of the main region was not correlated with survival, which serves as a negative control of the feature extraction process.

**Figure 3.**
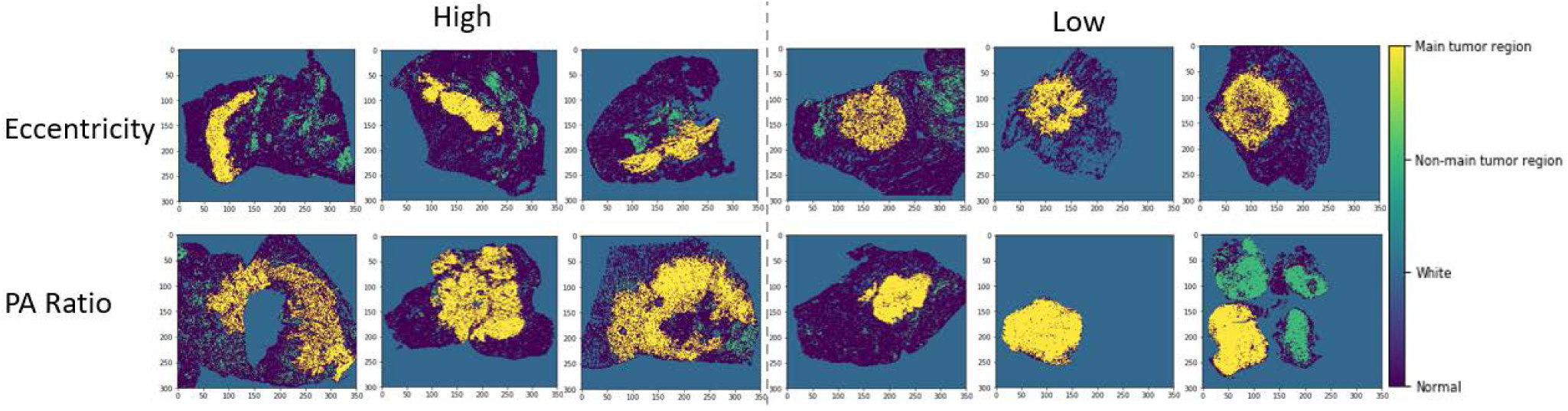
Comparison of tumor shapes with high or low values of eccentricity and PA ratio of main tumor region. Original heatmaps are cropped to the same size with the same image scale. Yellow, main tumor region; green, non-main tumor region; dark blue, non-malignant tissue; blue, blank part of pathology slide. P-A ratio, perimeter^2 to area ratio.

**Table 2.** Univariate analysis of tumor region features in NLST training dataset.

|  | *HR (95% CI)* | *p-value* |
| --- | --- | --- |
| *Number of regions (per 1000)* | 1.29 (0.64-2.58) | 0.48 |
| *Area sum of all regions (per 1000 pixels\*)* | 1.030 (1.010-1.050) | 0.0033 |
| *Perimeter sum of all regions (per 1000 pixels)* | 1.088 (1.028-1.151) | 0.0034 |
| *Sum of convex area for all regions (per 1000 pixels)* | 1.020 (1.006-1.033) | 0.0047 |
| *Sum of filled area for all regions (per 1000 pixels)* | 1.027 (1.009-1.045) | 0.0029 |
| *Sum of hole numbers of all regions (per 100)* | 1.087 (1.031-1.16) | 0.0033 |
| *Sum of major axis length of all regions (per 1000 pixels)* | 1.40 (1.00-1.96) | 0.051 |
| *Sum of minor axis length of all regions (per 1000 pixels)* | 2.65 (1.10-6.40) | 0.030 |
| *Perimeter<sup>2</sup>/area of all regions (per 1000)* | 1.18 (1.03-1.35) | 0.019 |
| *Area of main region (per 1000 pixels)* | 1.027 (1.007-1.048) | 0.0093 |
| *Convex area of main region (per 1000 pixels)* | 1.018 (1.004-1.032) | 0.010 |
| *Eccentricity of main region* | 6.37 (0.57-71.56) | 0.13 |
| *Hole number of main region (per 100)* | 1.087 (1.020-1.15) | 0.0060 |
| *Extent of main region* | 4.90 (0.19-126.30) | 0.34 |
| *Filled area for main region (per 1000 pixels)* | 1.025 (1.007-1.043) | 0.0072 |
| *Major axis length for main region (per 100 pixels)* | 1.57 (1.11-2.21) | 0.0099 |
| *Minor axis length for main region (per 100 pixels)* | 1.73 (1.05-2.83) | 0.031 |
| *Angle between the X-axis and the major axis of main region* | 0.98 (0.64-1.50) | 0.92 |
| *Perimeter of main region (per 1000 pixels)* | 1.087 (1.023-1.15) | 0.0068 |
| *Solidity of main region* | 7.24 (0.45-117.40) | 0.16 |
| *Average tumor probability of the main region (per 0.10)* | 1.11 (0.53-2.24) | 0.78 |
| *Perimeter<sup>2</sup>/area for main region (per 1000)* | 1.21 (1.03-1.42) | 0.021 |

NLST, the National Lung Screening Trial.
\* 1 pixel in heatmap = 1 patch in 40X pathological slide. Patch size: 300 pixels \* 300 pixels.

### Development and validation of prognostic model

Utilizing the tumor shape features extracted from the pathology images in the NLST dataset, we developed a prognostic model to predict patient survival outcome. The model was then independently validated in the TCGA cohort. Each patient was assigned into a predicted high-or low-risk group based on the extracted tumor shape features of the patient (see Methods Section). The survival curves for the predicted high-and low-risk groups are shown in Figure 4. The patients in the predicted high-risk group had significantly worse survival outcome than those in the predicted low-risk group (log rank test, p value = 0.0029). The multivariate analysis shows that the predicted risk groups independently predicted survival outcome (high-vs. low-risk, HR = 2.25, 95% CI 1.34-3.77, p value = 0.0022, Table 3) after adjusting for age, gender, smoking status and stage. This indicates the risk group defined by tumor shape features is an independent prognostic factor, in addition to other clinical variables.

**Figure 4.**
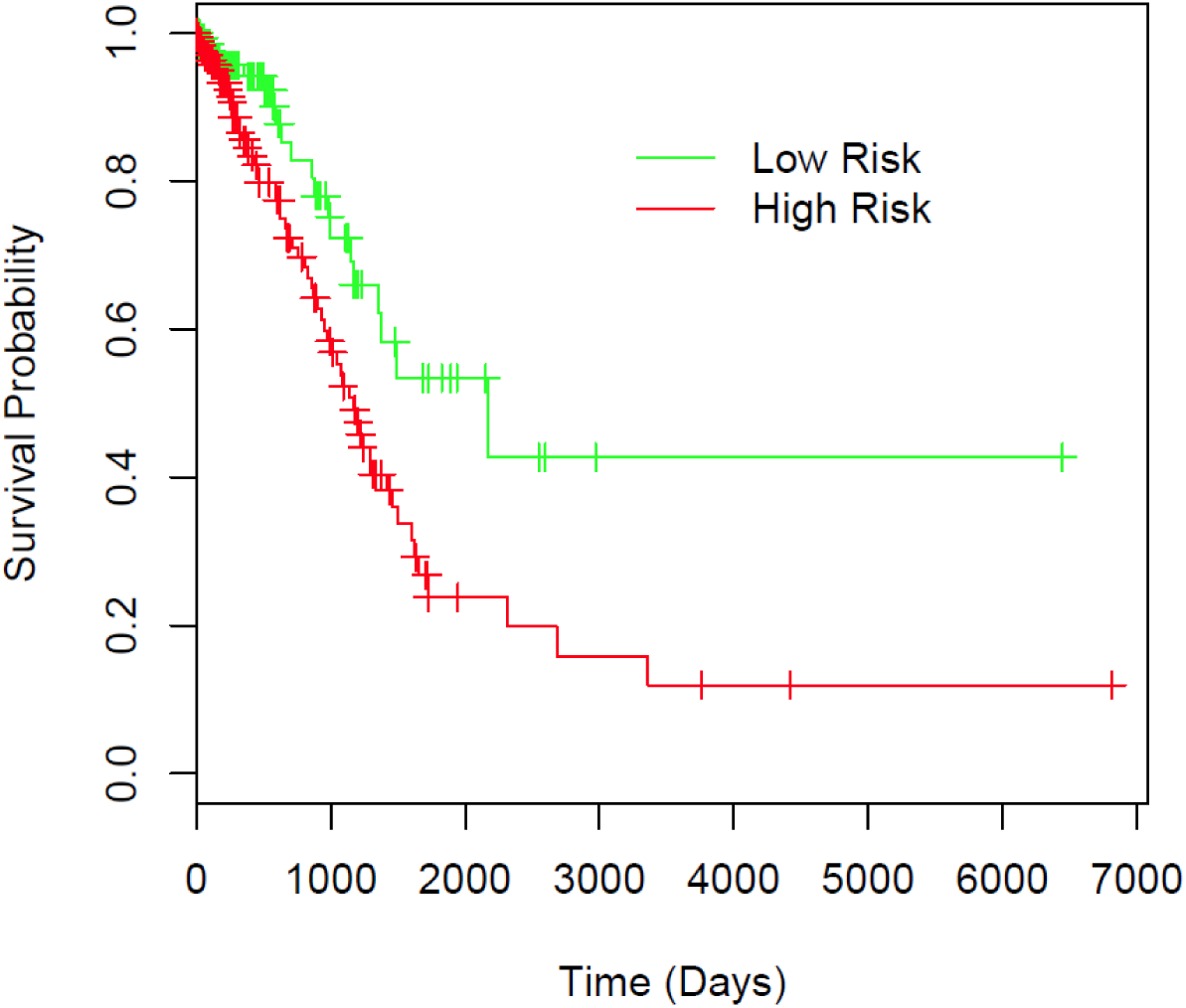
Prognostic performance in TCGA validation dataset illustrated by Kaplan-Meier plot. Patients are dichotomized according to median predicted risk score. Difference between the two risk groups: log-rank test, p value = 0.0029.

**Table 3.** Multivariate analysis of predicted risk and clinical variables in TCGA.

| *Variable* | *HR (95% CI)* | *p-value* |
| --- | --- | --- |
| *High risk vs. low risk* | 2.25 (1.34-3.77) | 0.0022 |
| *Age* | 1.02 (1.00-1.04) | 0.12 |
| *Male vs. female* | 0.71 (0.43-1.16) | 0.17 |
| *Smoker vs. non-smoker* | 0.95 (0.59-1.54) | 0.85 |
| *Stage II vs. stage I* | 2.58 (1.47-4.51) | <0.001 |
| *Stage III vs. stage I* | 5.23 (2.85-9.59) | <0.001 |
| *Stage IV vs. stage I* | 2.69 (1.19-6.09) | 0.017 |

## DISCUSSION

In this study, we developed image processing, tumor region recognition, image feature extraction, and risk-score prediction algorithms for pathology images of lung ADC. The algorithms successfully visualized the slide-level tumor region, and serve as a prognostic method independent of other clinical variables. The patient prognostic model was trained in the NLST cohort and independently validated in the TCGA cohort, which indicates the generalizability of the models to other lung ADC patient cohorts.

For tumor region detection, the whole slide image was divided into 300 × 300 pixel image patches, which were then classified into tumor, non-malignant, or white categories using a CNN model. The CNN model was trained on 3,848 images patches and tested on 1,068 patches, with 89.8% accuracy in the testing sets. Within the 109 incorrectly predicted patches, 27 contained an insufficient number of cells, which caused confusion between the tissue and background. For the other 82 cases where non-malignant patches were misclassified as tumor or vice versa, the cause seems to be interference from red blood cells, stroma, macrophages, and necrosis (Supplemental Table 1). The prediction errors related to out-of-focus tissues (such as macrophages and stroma cells) could be reduced by improved image scanning quality and training set labelling. A similar problem has also been reported in breast cancer recognition^14^.

The patch-level tumor prediction results were then arranged to generate tumor region heatmaps (Figure 2). In total, 22 well-defined image features were extracted for each tissue region, and averaged to generate patient-level features (Supplemental Figure 3). 15 of the 22 features were significantly correlated with survival outcome in NLST dataset (Table 2). The features related to tumor area and perimeter were associated with worse prognosis, as tumor size is a well-established prognostic factor, and size-based features were also reported as prognostic from lung cancer CT images^29,30^. Interestingly, both for all tumor regions and for the main tumor region, the perimeter^2 to area ratio was negatively correlated with survival outcome. The perimeter^2 to tumor area ratio is a quantification of the smoothness of the tumor surface; a large perimeter^2 to tumor area ratio indicates a large tumor surface and thus a rough tumor boundary. The negative correlation between perimeter^2 to area ratio and survival outcome is consistent with studies conducted on lung cancer CT images, which reported that a more irregular shape predicted worse survival^22,26^. To date, several genes have been reported to be associated with tumor shape^31,32^; understanding the relationship among gene expression, tumor shape, and survival outcome can provide insight into tumor development and guide therapeutic decision.

This is the first study to quantify tumor shape-related features using a CNN-based model in lung cancer. In addition, both the main tumor body and the tumor spread through air spaces (STAS, sometimes referred as aerogenous spread with floating cancer cell clusters [ASFC]), which can be easily detected in the heatmaps^33,34^. Since the median size of 40X pathology images is 24244 × 19261 pixels and the STASs usually only occupy 1 image patch (300 × 300 pixels) in the NLST dataset, it is labor intensive for human pathologists to circle accurate tumor boundaries and indicate all the tumor STASs. Thus, automatically generating the tumor region heatmap will facilitate pathologists in finding tumor regions and quantifying STASs. More importantly, our study has developed a computation-based method to quantify tumor shape, circularity, irregularity and surface smoothness, which can be an essential tool to study the underlying biological mechanisms. Although tumor size is a well-known prognostic factor, quantifications of the tumor area and perimeter-related features from pathology images are challenging and time-consuming for human pathologists. Thus, it is a natural step to extract image features directly from the predicted tumor heatmaps, thereby avoiding a subjective assessment by a human pathologist.

There are several limitations to our slide-level tumor region detection and image feature extraction pipeline. First, as mentioned before, our CNN model is sensitive to out-of-focus tissue such as red blood cells, macrophages, and stroma cells. Better pathology image scanning quality and more comprehensive labeling of the training set will help solve the problem. Second, the image features can be affected by slide preparation artifacts, such as artificially damaged tumor tissues and failure to select the slides that faithfully represent the tumor. Thus, to ensure the representability of the predicted risk score, a representative tumor slide is required. Third, the image slides are 2-dimensional, which loses the 3-dimensional spatial information. Combining the tumor prediction and feature extraction algorithms with other imaging techniques, such as CT or X-Ray, may produce more comprehensive descriptions of the tumor region and improve the performance of the current risk prediction model.

## CONCLUSION

Our pipeline for tumor region recognition and risk-score prediction based on tumor shape features serves as an objective prognostic method independent of other clinical variables, including age, gender, smoking status and stage. The tumor region heatmaps generated by our model can help pathologists locate tumor regions in gigapixel pathology slides swiftly and accurately. The model development pipeline can also be used in other cancer types, such as breast and kidney cancer.

## METHODS

### Datasets

The pathology images together with the corresponding clinical data were obtained from two independent datasets: 267 40X images for 150 lung ADC patients were acquired from the NLST dataset; 457 40X images for 389 lung ADC patients were acquired from the TCGA dataset. Clinical characteristics of patients in this study are presented in Table 1. The prognostic model was trained on the NLST dataset and independently validated on the TCGA dataset.

**Table 1.** Patient characteristics of training and validation datasets.

|  | *NLST (training)\** | *TCGA (validation)* | *p-value* |
| --- | --- | --- | --- |
| *No. of patients* | 150 | 389 |  |
| *Age* | 64.03 ± 5.12 | 64.98 ± 10.33 | 0.16 |
| *Gender* |  |  | 0.055 |
| *Male* | 82 (54.7) | 175 (45.0) |  |
| *Female* | 68 (45.3) | 214 (55.0) |  |
| *Smoking status* |  |  | 0.0020 |
| *Yes* | 81 (54.0) | 267 (68.6) |  |
| *No* | 69 (46.0) | 122 (31.4) |  |
| *Stage* |  |  | 0.0048 |
| *I* | 101 (67.3) | 222 (57.1) |  |
| *II* | 16 (10.7) | 96 (24.7) |  |
| *III* | 23 (15.3) | 49 (12.6) |  |
| *IV* | 10 (6.7) | 22 (5.7) |  |

NLST, the National Lung Screening Trial; TCGA, the Cancer Genome Atlas.
\* Values are either mean ± standard deviation, or number (percentage)

### Image patch generation

A CNN model was trained to classify non-malignant tissues, tumor tissues, and white regions based on image patches of H&E stained pathology images. The patch size was determined as 300 × 300 pixels under 40X magnification, to ensure at least 20 cells within one patch (Supplemental Figure 1). Tumor and non-malignant patches were randomly extracted from tumor regions and non-malignant regions labeled by a pathologist, respectively. The patches were classified as white if the mean intensity of all pixel values was larger than a threshold determined from sample images. Examples of each patch class are shown in Supplemental Figure 1. 2139 non-malignant, 2475 tumor and 730 white patches were generated in total.

### CNN training process

The Inception (V3) architecture^35^ with input size 300 × 300 and weights pre-trained on ImageNet was used to train our CNN model. The network was trained with stochastic gradient descent algorithms in Keras with TensorFlow backend^36^. The batch size was set to 32, the learning rate was set to 0.0001 without decay, and the momentum was set to 0.9. From the extracted 5,344 image patches, 3,848 patches (72%) were allocated to the training set, 428 patches (8%) to the validation set, and the remaining 1,068 patches (20%) to the testing set, with equal proportions among the three classes. Keras Image Generators were used to non-malignantize and flip the images, both horizontally and vertically, to augment the training and validation datasets. The maximum number of epochs to train was set to 50. To avoid overfitting, the training process automatically stopped after the validation accuracy failed to improve for 10 epochs.

### Prediction heatmap generation

To avoid prediction on a large empty image area and speed up the slide-level prediction process, the Otsu thresholding method followed by morphological operations such as dilation and erosion was first applied to whole pathology slides to generate the tissue region mask (Supplemental Figure 2)^37,38^. A 300 × 300 pixel window was then slid over the entire mask without overlapping between any two windows. The image patches were predicted with batch size 32, and one image patch was predicted only once without rotation or flipping. For each image patch, probabilities of being in each of the three classes were predicted, and a heatmap of the predicted probability was generated for each pathology slide (Figure 2). For each image patch, the class with the highest probability was determined as the predicted class.

### Image feature extraction

In a pathology slide, sometimes there are multiple tissue samples. To distinguish different tissue samples in the same slide, disconnected tissue regions were first identified by morphological operations on heatmaps of predicted classes (Supplemental Figure 3A-C)^38^. To remove the effects of some very small tissue samples, the tissue regions with area smaller than half of the largest tissue region in the same slide were removed from analysis. Within each tissue region, the tumor region with the largest area was regarded as the “main tumor region” (Supplemental Figure 3D). The following features of tumor regions were estimated for each tissue sample: number of regions, area, convex area, filled area, perimeter, major axis length, minor axis length, number of holes, and perimeter^2 to area ratio for all tumor regions and the main tumor region separately; eccentricity, extent, solidity, and angle between the X-axis and the major axis for the main tumor region (22 features in total)^39^. Here, 8-connectivity was used to determine disconnected tumor regions and disconnected holes^39^. When multiple tissue regions were available for one patient, either due to multiple tissues within one slide or multiple slides for one patient, the 22 image features were averaged to generate patient-level image features.

### Prognostic model development

A univariate Cox proportional hazard model was used to study the association between the 22 tumor shape features and patient survival outcome in the NLST dataset. The image features that were significantly associated with survival outcome were selected to build the prediction model for patient prognosis. To avoid overfitting, a Cox proportional hazard model with an elastic-net penalty^40^ was used; the penalty coefficient λ was determined through 10-fold cross-validation in the NLST cohort.

### Model validation in an independent cohort

The model developed from the NLST cohort was then independently validated in the TCGA cohort (n=389) for prognostic performance. First, the tumor region(s) in each pathology image slide from the TCGA dataset were detected using the CNN model, and the tumor shape features were extracted based on the detected tumor region(s) in each image slide and summarized to patient level. Finally a risk score was calculated based on the extracted tumor features. The patients were dichotomized into high-and low-risk groups based on their predicted risk score, using the median as the cutoff. A log-rank test was used to compare survival difference between predicted high-and low-risk groups. The survival curves were estimated using the Kaplan-Meier method. A multivariate Cox proportional hazard model was used to test the prognostic value of the predicted risk score after adjusting for other clinical factors, including age, gender, tobacco history and stage. Overall survival, defined as the period from diagnosis until death or last contact, was used as response. Survival analysis was performed with R software, version 3.3.2^41^. R packages “survival” (version 2.40-1) and “glmnet” (version 2.0-5) are used^40,42^. The results were considered significant if the two-tailed p value ≤ 0.05.

## Data availability

Pathology images that support the findings of this study are available online in NLST (https://biometry.nci.nih.gov/cdas/nlst/) and The Cancer Genome Atlas Lung Adenocarcinoma (TCGA-LUAD, https://wiki.cancerimagingarchive.net/display/Public/TCGA-LUAD).

## Code availability

The codes are available upon request. We will share the codes through Github following manuscript acceptance.

## Author Contributions

SW and GX designed the study. SW, AC, LC, YX, JF, AG, and GX analyzed the results and wrote the manuscript. SW and AC implemented the algorithm. LY labeled the data. All authors commented on the manuscript.

## Materials & Correspondence

Correspondence and material requests should be addressed to GX.

## Supplementary material

**Supplemental Figure 1.**
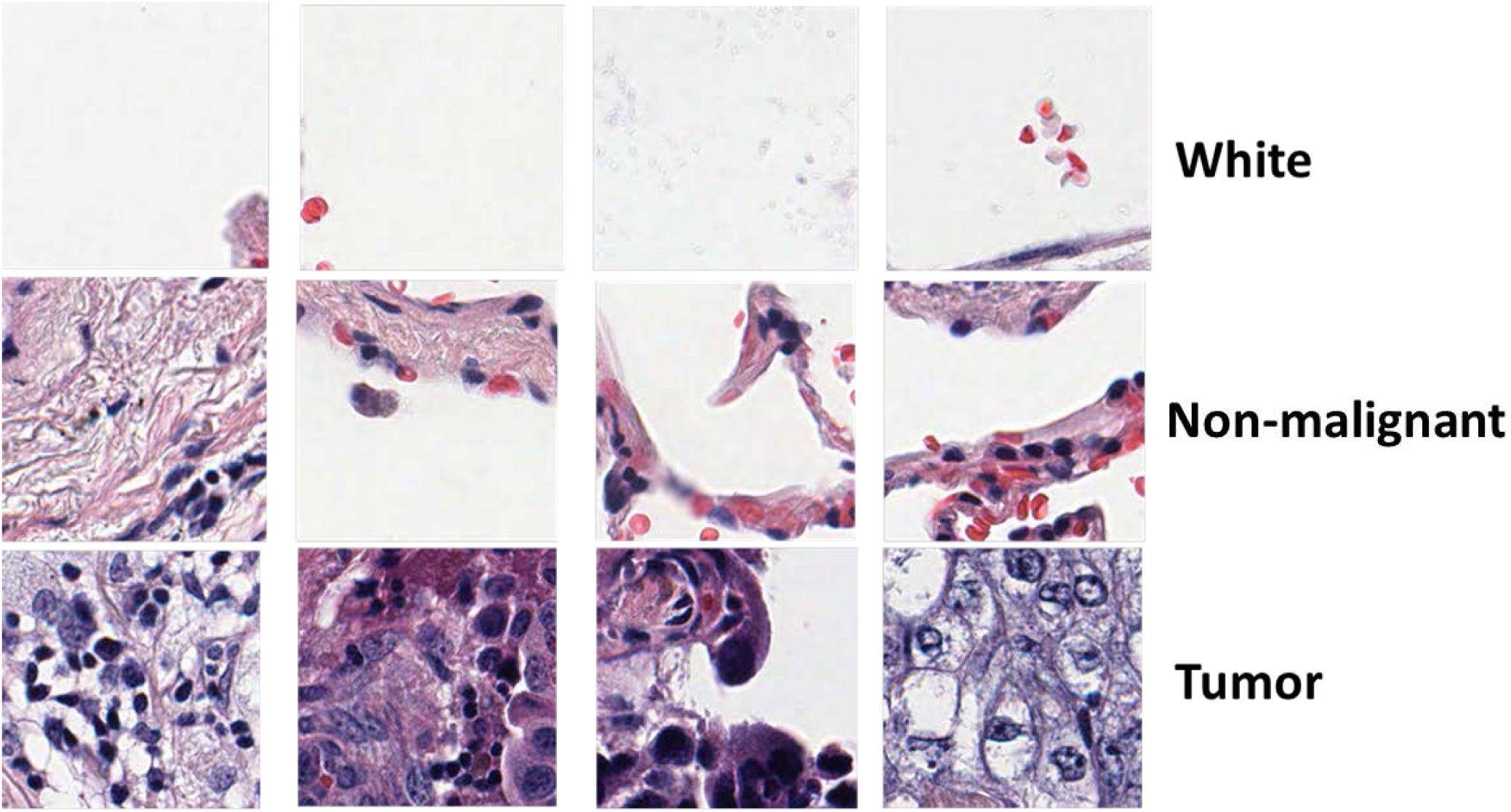
Example of image patches from “white” (empty regions, upper panel), “non-malignant” (middle panel) and “tumor” (bottom panel) categories. Patch size: 300 × 300 pixels.

**Supplemental Figure 2.**
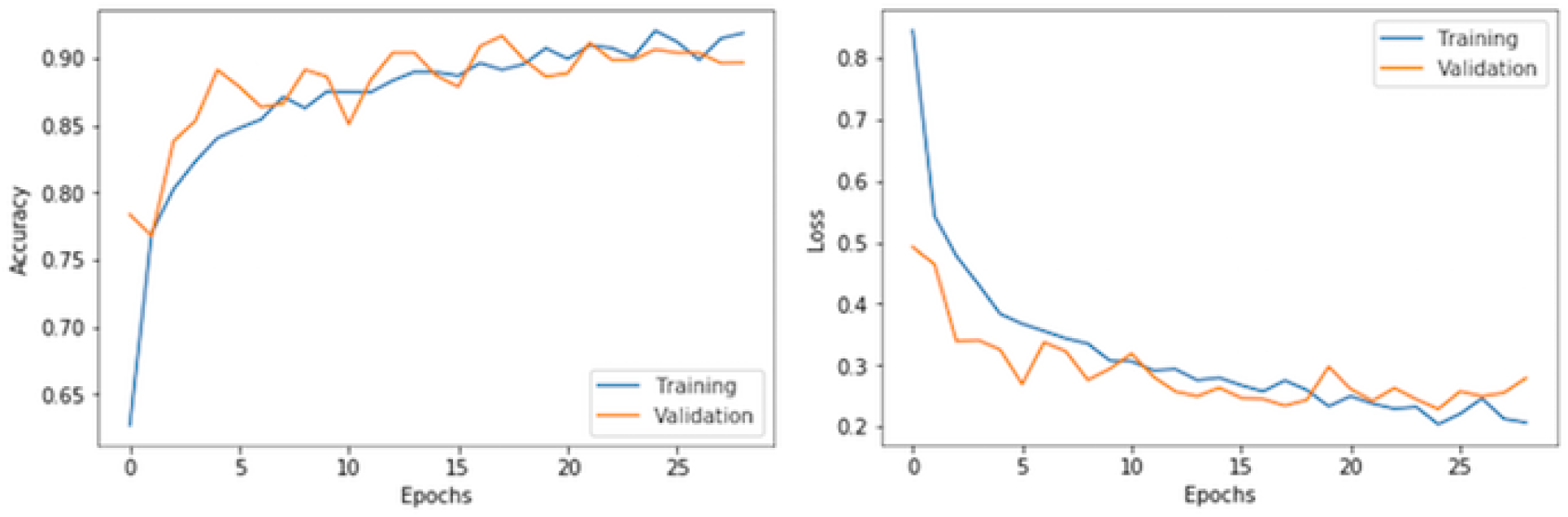
Convolutional Neural Network learning curves in both training and validation datasets. Left, accuracy versus epochs; right, loss versus epochs.

**Supplemental Figure 3.**
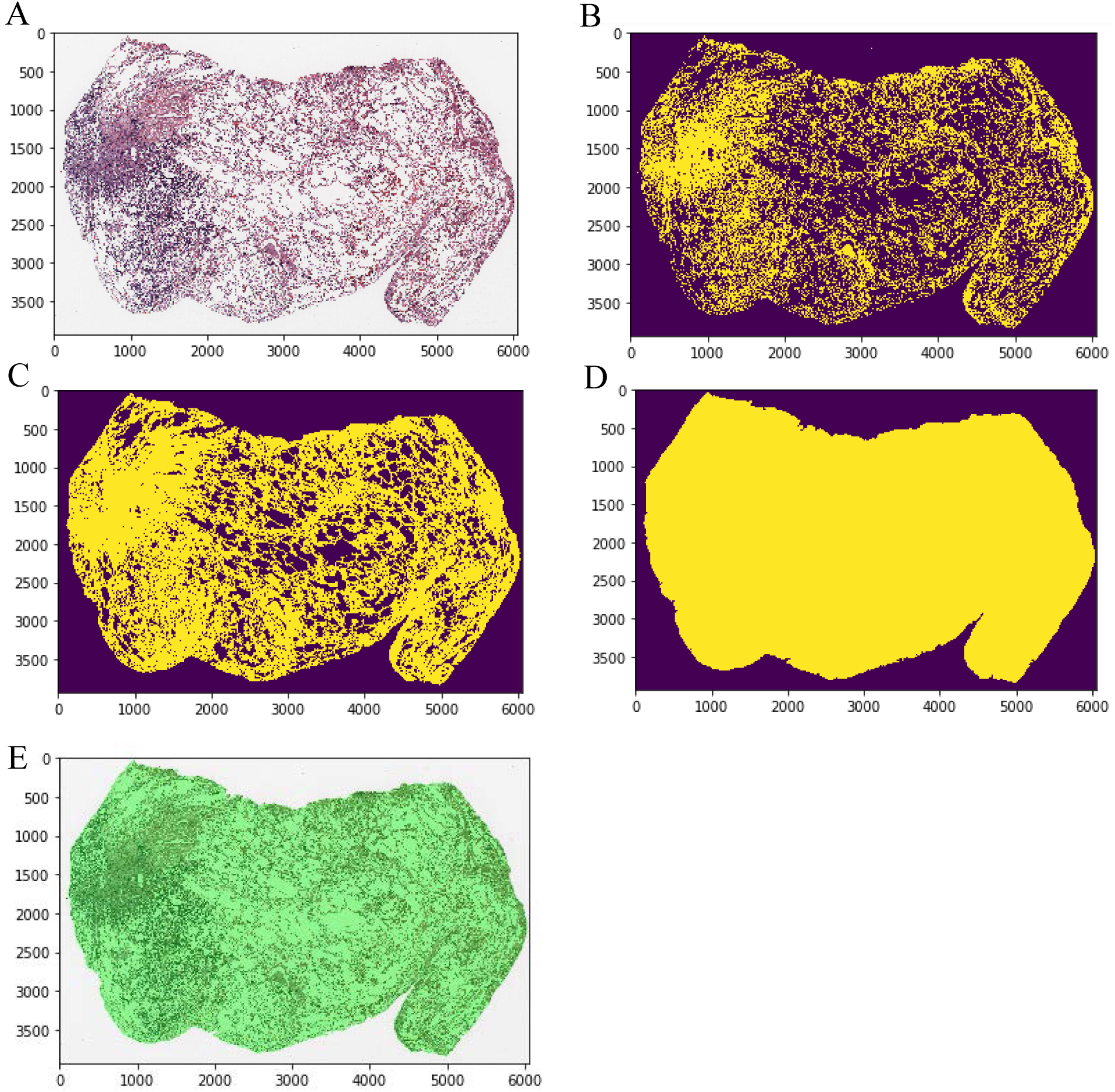
Otsu thresholding and image morphological operations to speed up slide level prediction process. (A) The original slide. (B) The image mask after Otsu thresholding. (C) The image mask after dilation and removing small objects of the mask in (B). (D) The final mask after dilation, erosion, and filling up holes of mask in (C). (E) Overlap final image mask and original pathology slide.

**Supplemental Figure 4.**
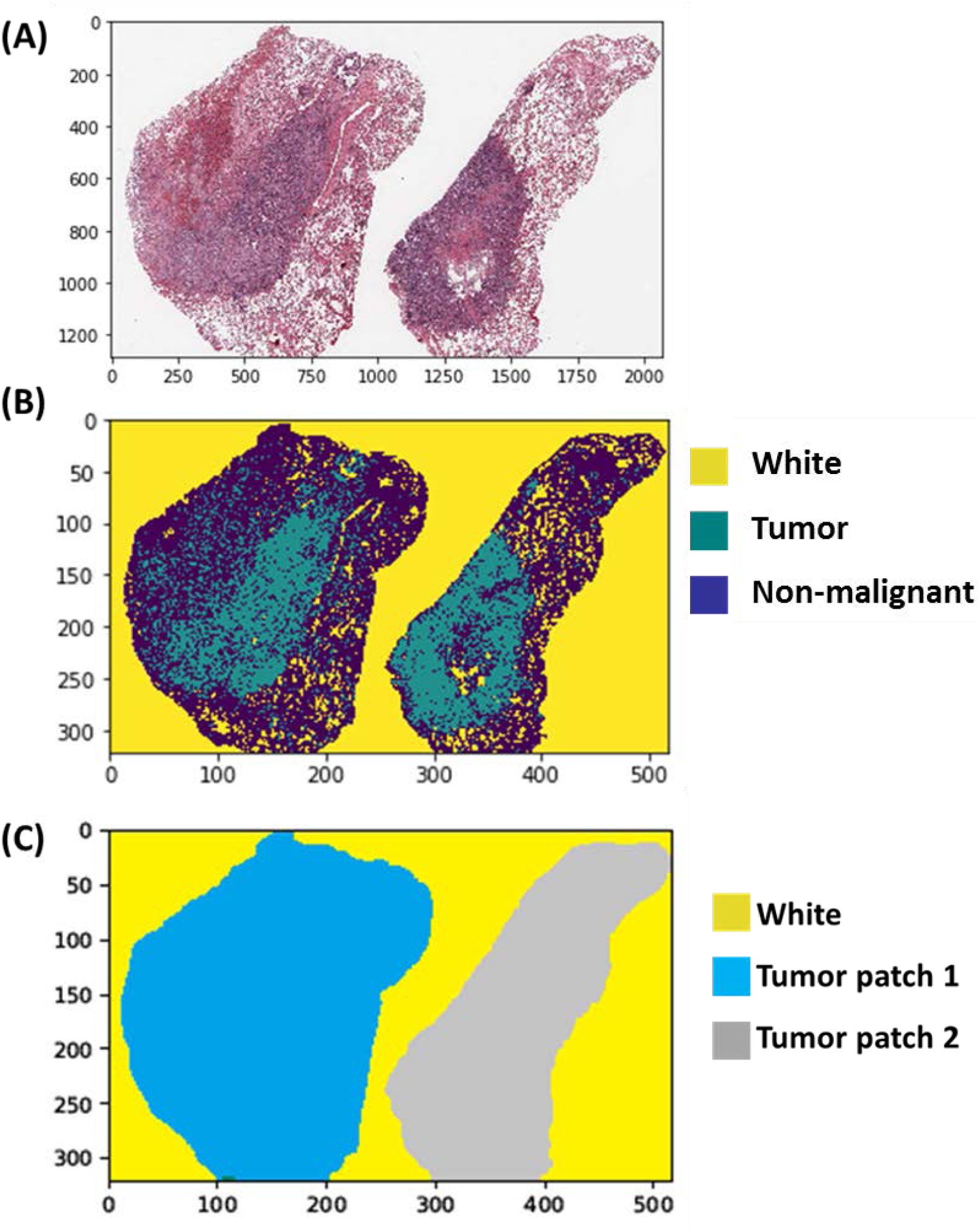
Tissue region identification in case of multiple tissue samples within one slide. **(A)** Original slide. **(B)** Predicted patch-level tumor, non-malignant and white heatmap. **(C)** Disconnected tissue samples identified by image processing. Yellow, background; blue, first tissue patch; gray, second tissue patch.

**Supplemental Table 1.** Confusion matrix for image patch classification.

| *Ground Truth \ Predicted Value* | *Non-malignant* | *Tumor* | *White* |
| --- | --- | --- | --- |
| *Non-malignant* | 400 (93.5%) | 24 (5.6%) | 4 (0.9%) |
| *Tumor* | 58 (11.7%) | 436 (88.1%) | 1 (0.2%) |
| *White* | 21 (14.4%) | 1 (0.7%) | 124 (84.9%) |

**Supplemental Table 2.** Comparison of patient characteristics between high-risk and low-risk groups in TCGA validation dataset.

|  | *Low-risk* | *High-risk* | *p-value* |
| --- | --- | --- | --- |
| *No. of patients* | 195 | 194 |  |
| *Age* | 65.50 ± 10.11 | 64.45 ± 10.54 | 0.31 |
| *Gender* |  |  | 0.20 |
| *Male* | 81 (41.5) | 94 (48.5) |  |
| *Female* | 114 (58.5) | 100 (51.5) |  |
| *Smoking status* |  |  | 0.89 |
| *Yes* | 135 (69.2) | 132 (68.0) |  |
| *No* | 60 (30.8) | 62 (32.0) |  |
| *Stage* |  |  | 0.005 |
| *I* | 127 (65.1) | 95 (49.0) |  |
| *II* | 44 (22.6) | 52 (26.8) |  |
| *III* | 17 (8.7) | 32 (16.5) |  |
| *IV* | 7 (3.6) | 15 (7.7) |  |

